# Assessment of label-free quantification and missing value imputation for proteomics in non-human primates

**DOI:** 10.1101/2021.07.30.454221

**Authors:** Zeeshan Hamid, Kip D. Zimmerman, Hector Guillen-Ahlers, Cun Li, Peter Nathanielsz, Laura A. Cox, Michael Olivier

**Affiliations:** Center for Precision Medicine, Wake Forest University School of Medicine, Winston-Salem, NC; Southwest National Primate Research Center, San Antonio, TX; Department of Animal Science, University of Wyoming, Laramie, WY

**Author notes:** **Correspondence to:** Michael Olivier, PhD, Center for Precision Medicine, Nutrition Research Center, G-floor, Wake Forest School of Medicine, Medical Center Blvd, Winston-Salem, NC 27157.

**Keywords:** Non-human primates (NHP), Posttranslational modifications (PTMs), Missing Value imputation

## Abstract

**Introduction:** Reliable and effective label-free quantification (LFQ) analyses are dependent not only on the method of data acquisition in the mass spectrometer, but also on the downstream data processing, including software tools, query database, data normalization and imputation. In non-human primates (NHP), LFQ is challenging because the query databases for NHP are limited since the genomes of these species are not comprehensively annotated. This invariably results in limited discovery of proteins and associated Post Translational Modifications (PTMs) and a higher fraction of missing data points. While identification of fewer proteins and PTMs due to database limitations can negatively impact uncovering important and meaningful biological information, missing data also limits downstream analyses (e.g., multivariate analyses), decreases statistical power, biases statistical inference, and makes biological interpretation of the data more challenging. In this study we attempted to address both issues: first, we used the MetaMorphues proteomics search engine to counter the limits of NHP query databases and maximize the discovery of proteins and associated PTMs, and second, we evaluated different imputation methods for accurate data inference.

**Results:** Using the MetaMorpheus proteomics search engine we obtained quantitative data for 1,622 proteins and 10,634 peptides including 58 different PTMs (biological, metal and artifacts) across a diverse age range of NHP brain frontal cortex. However, among the 1,622 proteins identified, only 293 proteins were quantified across all samples with no missing values, emphasizing the importance of implementing an accurate and statiscaly valid imputation method to fill in missing data. In our imputation analysis we demonstrate that Single Imputation methods that borrow information from correlated proteins such as Generalized Ridge Regression (GRR), Random Forest (RF), local least squares (LLS), and a Bayesian Principal Component Analysis methods (BPCA), are able to estimate missing protein abundance values with great accuracy.

**Conclusions:** Overall, this study offers a detailed comparative analysis of LFQ data generated in NHP and proposes strategies for improved LFQ in NHP proteomics data.

## Introduction

Quantitative proteomics using mass spectrometry (MS) has significantly advanced over the years and now allows the effective analysis of thousands of proteins. A number of technical advancements have been made at both the instrument level and data analysis level to increase the amount of biologically relevant information elicited from proteomics data. One of the methods for mass spectrometry-based proteomics is label-free quantification (LFQ) where the relative amount of proteins across experimental groups is determined without using any isotopically labelled tags [1, 2]. LFQ proteomics has several advantages for large scale experiments in terms of cost effectiveness and overall experimental design. Also, with the advent of high resolution top end mass spectrometers, LFQ has become more reliable and a method of choice for many investigators [3].

A primary challenge for proteomics data analysis is the direct dependence on predefined protein sequence databases for protein identification. During the proteomics data search, experimental data are searched against an in silico-generated library of theoretical spectra obtained from the gene annotation of the same organism. Therefore, when the gene annotation of the organism is not very detailed, it directly and negatively impacts the resulting proteomics data. Specifically, reduced genome annotation results in fewer protein identifications and inadequate detection of important PTMs. Genetically well annotated organisms like *H. sapiens or M. musculus* benefit from rich genome annotations, allowing robust proteomics data searches that account for known genetic variability like splice variants or amino acid sequence variation of encoded proteins [4]. For proteomics in other organisms, a different approach is needed to account for the incomplete gene annotation and less robust proteomics data search. To this end, our group has previously shown that the use of proteogenomic approaches can improve the search library limitations by incorporating the information from transcriptomics data into search databases [4]. However, adapting proteogenomic approaches can be cumbersome, time-consuming, and cost-intensive since it requires additional RNA-Seq experiments, and in some cases proteogenomic approaches may be impossible due to limited amounts of available samples.

Another challenge in proteomics, particularly for LFQ, is the occurance of missing data. Missing data in LFQ can range from 10-50% overall, and in 70-90% of data across multiple samples at least 1 data point is missing per peptide/protein [5]. The sources of missing values range from tryptic miscleavages to ion suppression in the mass spectrometer, and improper MS/MS fragmentation [5]. Because LFQ data contain a relatively high percentage of missing values, multiple approaches for the imputation of missing values in proteomics data have been proposed [5-8]. However, there is no unified consensus on a best approach for imputing missing values in proteomics data and there has been little discussion about applying Multiple Imputation (MI) methods [5-10]. MI methods are often implemented to account for the uncertainty in the prediction of the imputed values, whereas Single Imputation (SI) methods treat the predicted values as if they were true values in downstream association analysis. MI methods reduce bias in statistical analysis by creating multiple versions of the imputed data where the imputed values in each of the imputed datasets are allowed to vary by a random component that reflects the level of confidence the prediction algorithm has in a particular imputed value [11]. In the statistical literature, MI is preferred over SI because SI can falsely inflate correlations between variables and lead to biased statistical inference by treating imputed values as if they were true values in downstream association analysis [11-15].

In this study, we tested the effectiveness of various approaches to improve both the proteomics search coverage and enhance quantitative data information. We first implemented the MetaMorphues proteomics search engine which can enhance proteomics or PTM searches and compensate to a greater extent for less well annotated genomes and search databases [16]. Then we evaluated different imputation approaches for filling in missing data points, comparing SI and MI methods. Only comparing the accuracy of the raw imputed values between SI and MI methods is not appropriate, because the imputed raw values from MI methods will always appear less accurate by design [11-15]. As a more appropriate way of comparing the two techniques, we first compared the accuracy of regression coefficients from downstream association analyses computed on the incomplete data imputed by SI methods with the regression coefficients computed by association analysis on the real and complete data. Secondly, we compared the pooled regression coefficients from MI with regression coefficients computed by association analysis on the real and complete data. In the context of high-dimensional proteomics data, MI techniques are not as well explored [9, 10, 17]. Here, we examined how well MI imputes high-dimensional data and whether or not MI is necessary to reduce bias when imputing missing values with proteomics data for association analysis. A short overview of our study design is shown in Fig. 1. Overall, our study provides a detailed description for an improved pipeline for label free proteomics in non-human primates (NHP).

**Fig. 1.**
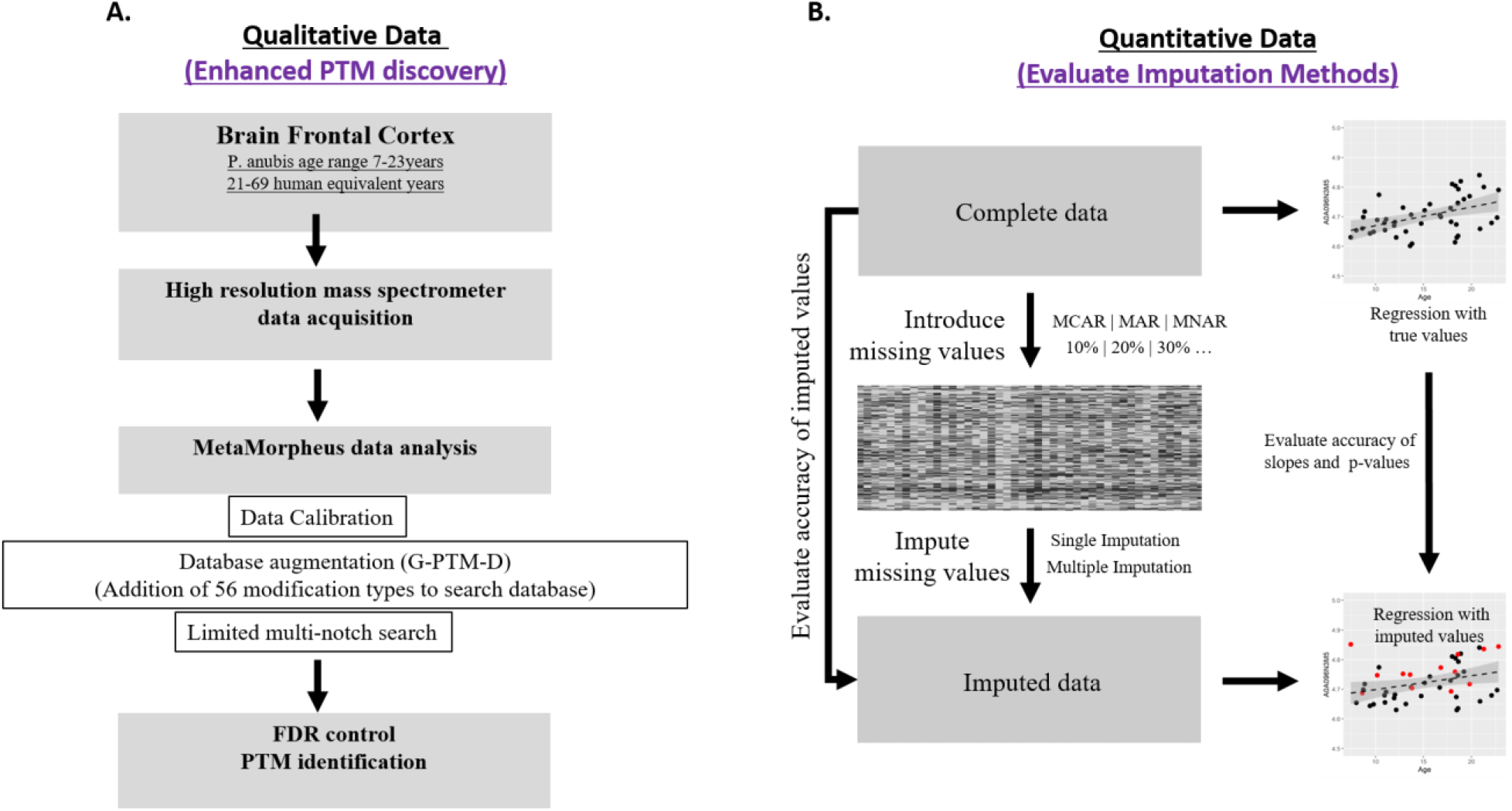
Overview of the study design **(A)** Enhanced discovery of PTMs using high resolution mass spectrometry and search database augmentation in MetaMorphues **(B)** Graphical representation of imputation workflow (missing data introduction and comparative evaluation of single and multiple imputation).

## Results and Discussion

The raw proteomics data search using MetaMorpheus identified 1,622 proteins and 10,634 peptides across all 45 *P. anubis* prefrontal cortex samples (Age 7-23 years). As described in a study by Solntsev *et al*., MetaMorpheus allows faster and more accurate proteomics data searches with comprehensive analysis of PTMs using a G-PTM-D discovery approach. It uses a multi-notch search, a hybrid search strategy between traditional narrow window search and open search. In the multi-notch strategy, the search is narrowed compared to an open search to only allow certain predefined mass differences for peptide modifications, thus decreasing the search time. Also, with the data calibration in MetaMorpheus there is increased accuracy of peptide identifications [16]. The modifications identified in our study include common biological PTMs (such as acetylation, oxidation, methylation or phosphorylation), common artifacts (carbamyl, deamidation, water loss or ammonia loss) and some metal PTMs (calcium, sodium, iron etc.). An overview of the observed peptides with different modifications is shown in Fig. 2. As shown in the upper panel of Fig. 2, out of 4,963 high confident peptides (q≤0.01, precursor mass within ±5ppm error tolerance, presence of at-least one unambiguous PTM and a MetaMorpheus score of >8), we found 3,254 peptides with no or fixed modifications only, 945 biological PTMs, 434 metal PTMs and 711 common artifacts (usually introduced during sample preparation or mass spectrometer data acquisition). Further breakdown of the number of peptides specific to each modification type is shown in the lower panel of Fig. 2. Our data showing oxidation at Methionine (M) and N-terminal acetylation being most common PTMs are in line with the literature and are therefore often included as variable modifications in proteomics search engines. Amino acid specificity and associated PTMs identified in our study are shown in Table 1. The most common biological modifications occur at Lysine (K) apart from other important biological modifications like ADP-Ribosylation, HexNac, phosphorylation, nitrosylation occurring at serine (S), threonine (T) and tyrosine (Y). Most of the metal-specific PTMs occur at aspartate and glutamate. The complete list of all normalized protein data and all peptide data is shown in Supp. File 1, and a list of observed PTM specific peptides is shown in Supp. File 2.

**Table 1:**
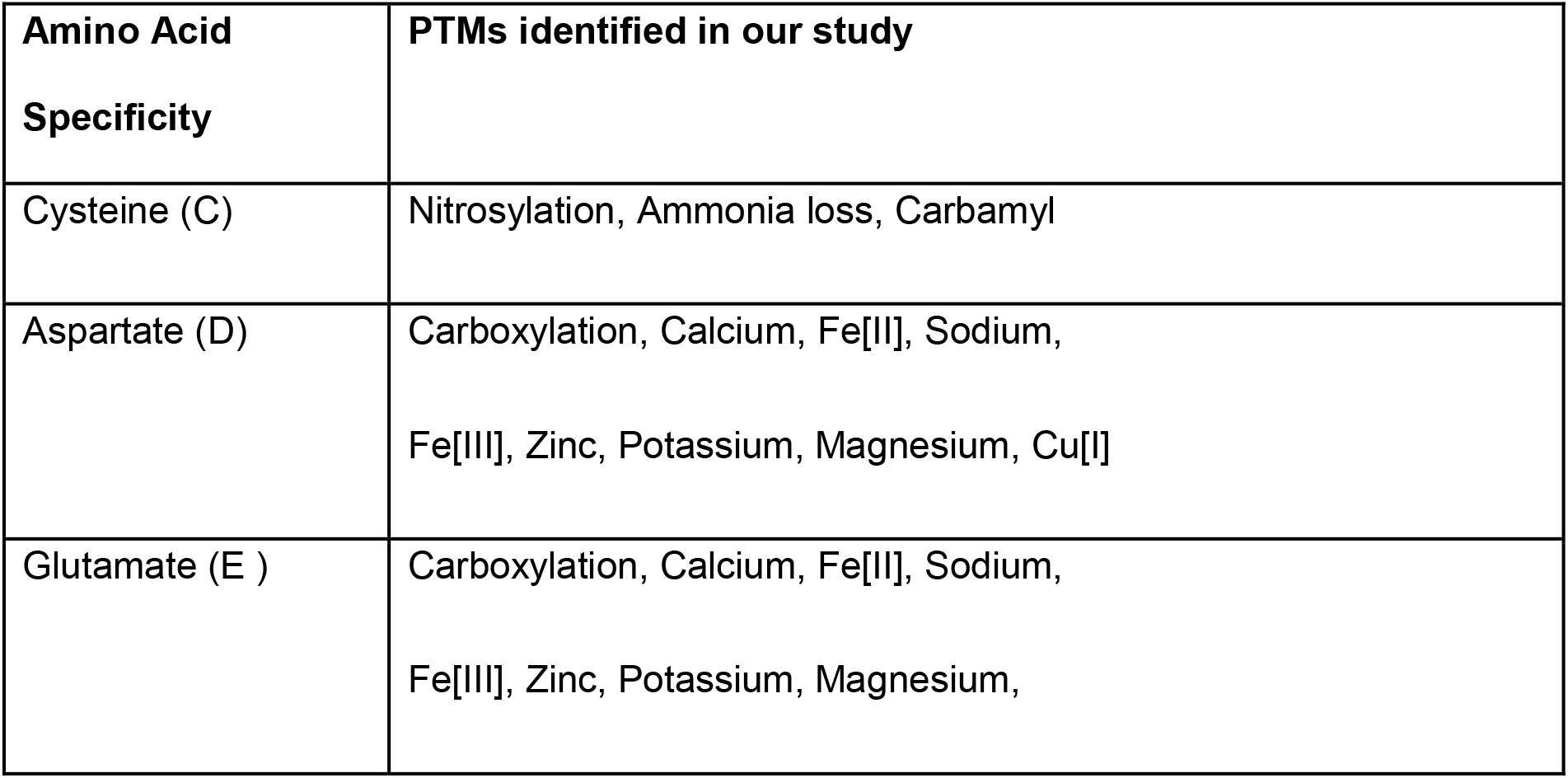
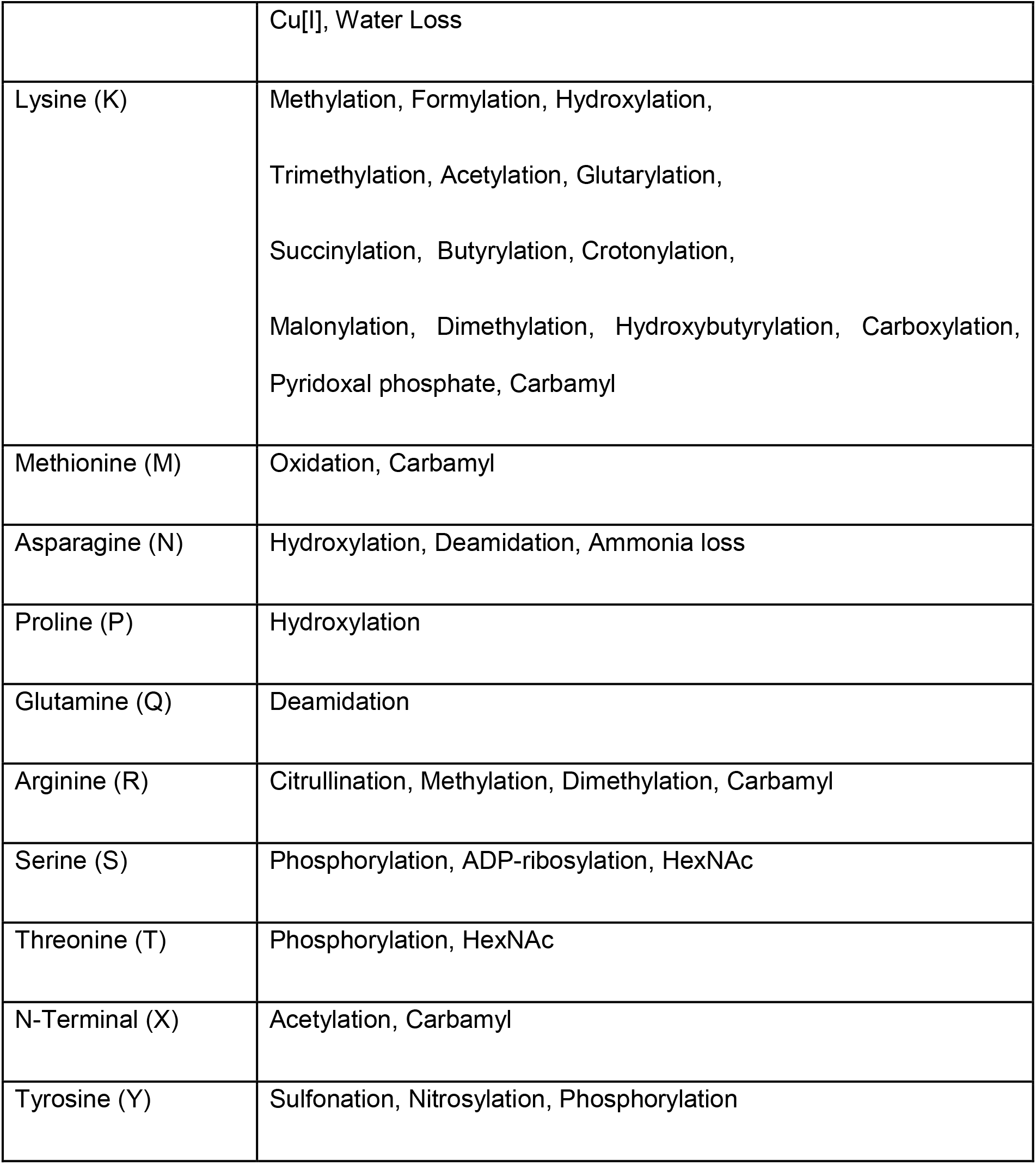
Amino acid specificity of detected PTMs in our study.

**Fig. 2.**
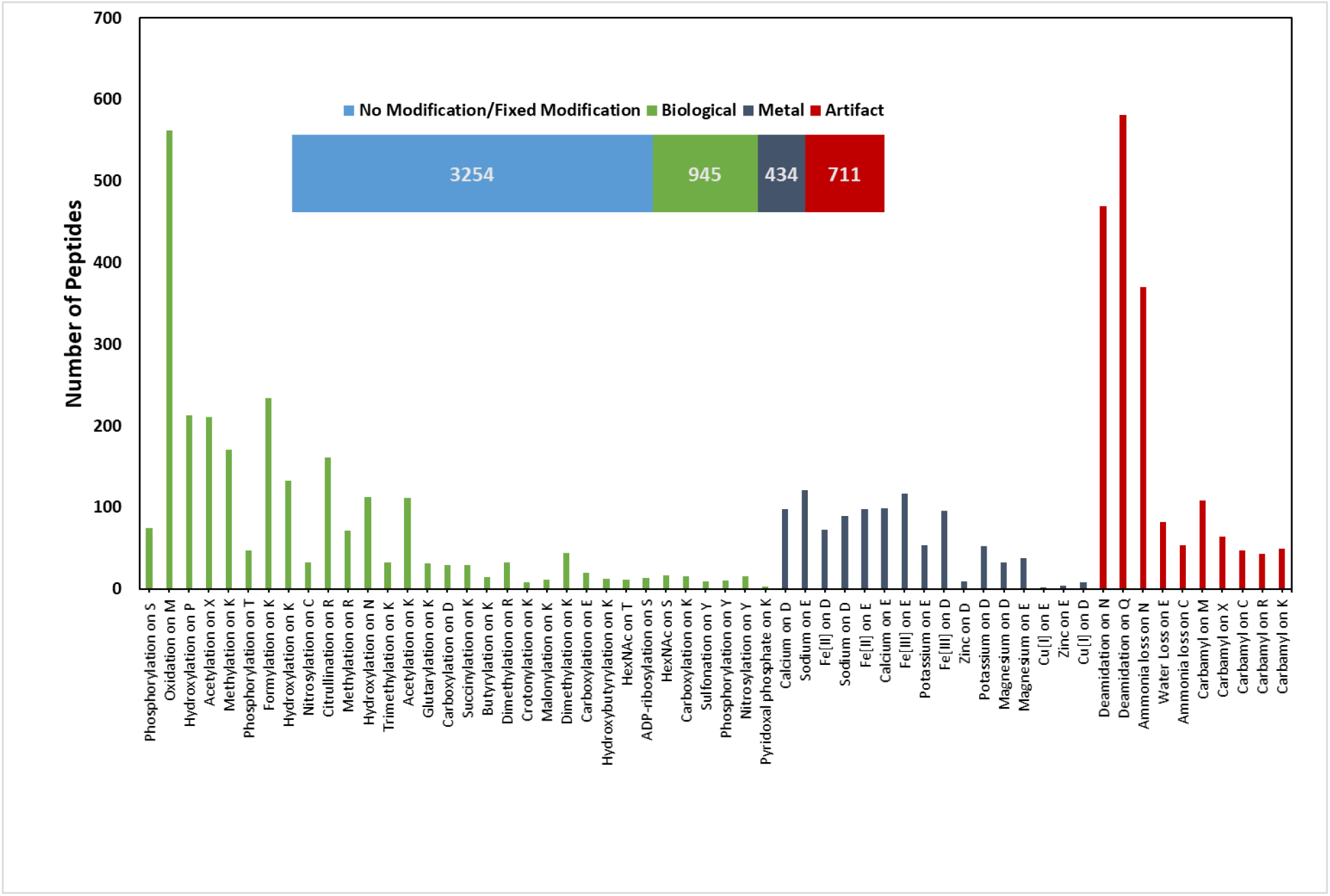
Distribution of peptides with or without modification (number in bars represent number of peptides observed in each category). Lower panel shows breakdown of number of PTM specific peptides detected in each subclass of three categories (Biological, Metal and Artifact)

To illustrate the importance of our PTM discovery in *P. anubis*, we selected Carbonic anhydrase 3 (CA3) as a reference protein. This protein is member of the carbonic anhydrase (CA) family with an important physiological role in hydration of carbon dioxide to bicarbonate and protons. Many CAs have been shown to be overexpressed in several diseases and are intriguing drug targets with some already in clinical use as therapeutic agents for glaucoma and epilepsy [18]. Using proteomics we can have a more in-depth understanding of several isoforms, along with their novel PTMs, which can provide additional leads in future drug discovery programs of CAs. With our current proteomics workflow, as shown in Supp. File 4 Panel A, we observed more than 90% of sequence coverage of CA3. Apart from some known modifications like phosphorylation at S48 (reported earlier in *Rattus norvegicus* UniProtKB - P14141 (CAH3_RAT) [19], we also observed a number of important and previously unknown modifications including various metal PTMs which might have important biological roles given that CA3 is part of the metalloprotein superfamily. Further studies are needed to confirm the biological importance of these novel modifications. A full list of all individual peptides and PTMs detected for CA3 are reported in Supp. File 3. For representative and manual confirmation of the PTM identification workflow, Supp. File 4 panel B shows all PTMs detected for reference peptide “DIKHDPSLQPWSVSYDGGSAK” from CA3 along with the experimental MS/MS spectral match with theoretical “b” and “y” ions.

Another independent but related focus of our study was to improve the quantitative accuracy of the proteomics data by imputing the missing data points. As the nature of the missing values is directly affected by the search algorithms, the availability of less comprehensive databases for our current model organism (*P. anubis* Uniprot Reference Proteome ID UP000028761) resulted in more missing data points. One of the approaches used for increasing proteomics data information and decreasing percent missing values for organisms having a less comprehensive database is by searching the data with combined search database using genetically related organisms with more comprehensive databases [20]. For our dataset, there is no significant increase in either the number of proteins or percentage of missing data when applying this approach (e.g. by using *H. sapiens* search database together with *P. anubis* we observed 1974 proteins, where 630 proteins were observed with *P. anubis* reference database only, 634 proteins were observed with *H. sapiens* reference database and 710 proteins were observed as shared between *P. anbuis* and *H. sapiens)*.

Even though MetaMorpheus (with its G-PTM-D and Match between runs option) compensates to a greater extent for the missing data arising due to less comprehensive search database for *P. anubis* and missing-ness arising due to other factors including improper MS/MS fragmentation, our protein data still contained a large percentage of missing data. Out of 1,622 total proteins identified, only 293 proteins had no missing values across all samples. This high number of proteins with incomplete data emphasizes the importance of implementing an accurate and statistically valid imputation method to fill in missing data. Otherwise, these missing data will decrease statistical power, introduce potential biases, and limit options for downstream analyses. With a limited availability of studies testing effectiveness of imputation strategies in NHP proteomics data, we aimed to evaluate various imputation methods on our current dataset to determine which method would be most helpful improving the statistical analysis of LFQ proteomics data for organisms with less comprehensive annotated genomes in general and NHP species in particular. Brief overviews of each of the various imputation methods used in this study are provided in Supp. File 5.

First we evaluated the accuracy of SI techniques. SI methods fill in the best predicted value for each missing data point and treat it as truth for the subsequent statistical analyses. We used several different SI methods in this study, including single value approaches where missing values are very simply replaced with a constant or a randomly selected value. Here we chose to only apply two single value approaches. The first method replaces missing values with zeros [5] (zero imputation) and the second replaces missing values with the respective protein mean (mean imputation). We also implemented approaches where information from the whole dataset is used to reconstruct the missing values (e.g., SVD, BPCA) [21-23]. Lastly, we applied methods that exploit measures of local similarity for imputation (e.g., KNN, GRR, and Random Forest) [22-25]. All of the approaches, however, are SI methods that do not incorporate uncertainty into the prediction of the imputed values.

In contrast, MI methods are implemented to account for the uncertainty in the prediction of the imputed values by creating multiple versions of the imputed data where the imputed values in each of the imputed datasets are allowed to vary by a random component that reflects the level of confidence the prediction algorithm has in a particular imputed value [11-15, 26]. It has been repeatedly demonstrated that MI methods can reduce bias in downstream statistical analysis [11-15]. For this reason, we evaluated MI techniques alongside SI methods and we specifically considered if any of the imputation methods were significantly altering correlations between samples and measures or causing biased statistical inference.

To evaluate the accuracy of the SI methods, we first introduced missing values into our complete data with varying degrees of global missingness (Supp. Fig. 1). Then, we imputed the missing data with each imputation method and compared the results to the original complete data. Percent bias (i.e., the percent difference between the imputed value and truth) was calculated for each method, and data points with a percent bias <5% were considered as correctly imputed. Fig. 3 shows the percentage of correctly imputed values using different methods for a mixture of MCAR-MAR-MNAR missingness. Our results indicate that SI methods exploiting local similarity (GRR, LLS, SeqKNN, IMPSeq, and Random Forest) or the global data structure (BPCA) all accurately impute the raw data at high proportions. Methods where missing values were replaced with the protein mean also showed reasonably high accuracy, but repeatedly filling in the mean for each missing value - particularly where there is high amounts of missingness - will cause distributional problems and will very likely introduce biases for statistical correlation analyses of the imputed data. We note that the SeqKNN and IMPSeq methods often failed to compute with our data in our implementation. There was not one SI method that demonstrated clear improvements in accuracy over the others, so we selected four methods (GRR, LLS, BPCA, and Random Forest) to be further evaluated at overall missingness percentages of 10-30% and protein missingness percentages between 0-30% (Supp. Fig. 2). Again, among these top four methods there was no significant difference in accuracy (ANOVA p = 0.19 for 30% global missingness and 20-30% protein missingness).

**Fig. 3.**
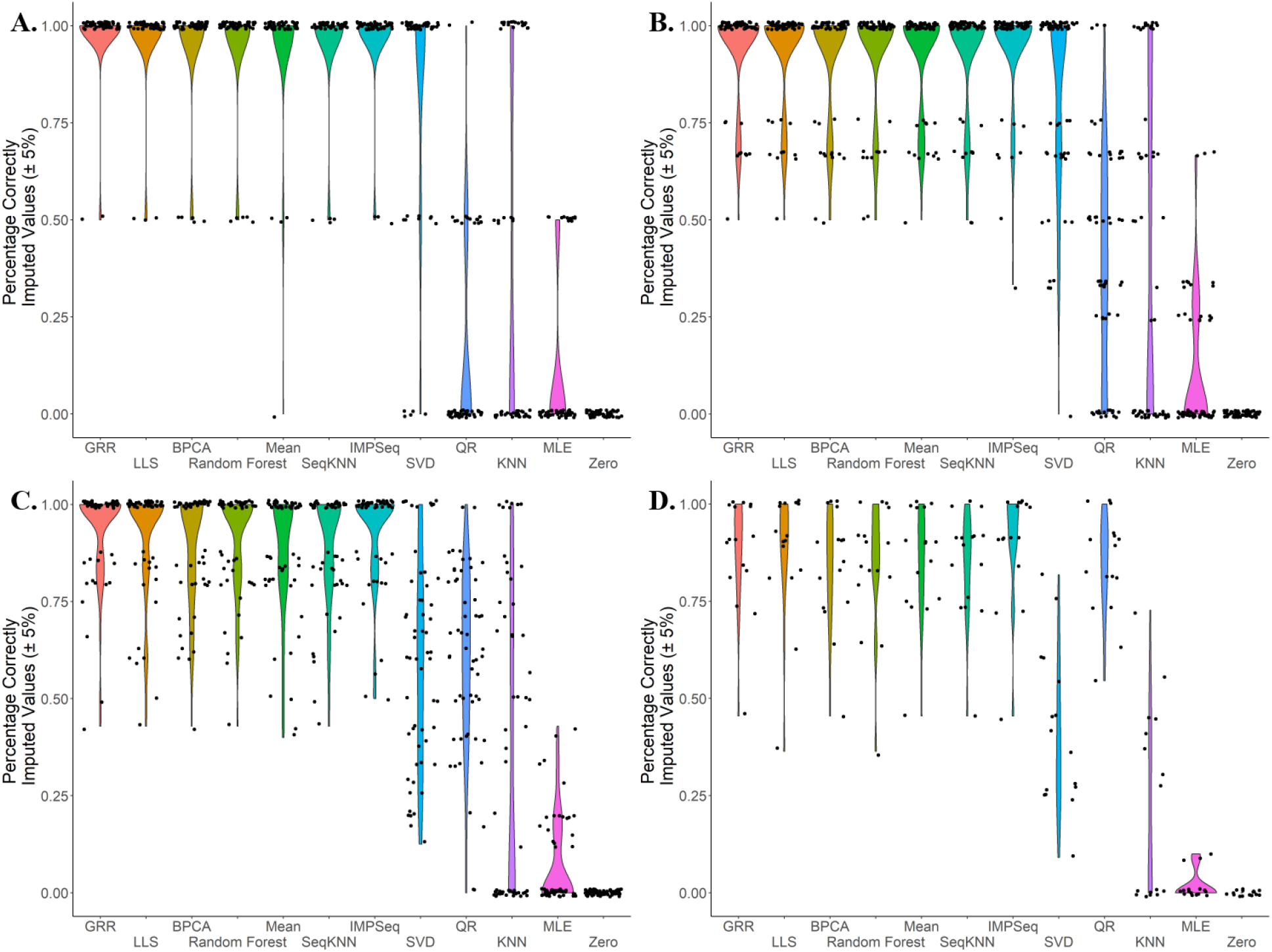
Percentage of correctly imputed values (imputed values with Percent Bias < 5%) for 12 different Single Imputation methods and 4 different levels of overall missingness (A. < 5% protein missingness B. 5-10% protein missingness C. 10-20% protein missingness D. 20-30% protein missingness.

Using the four highly accurate SI methods and two additional MI methods, we evaluated whether or not any of the methods introduced significant bias into the downstream statistical analysis. The data here are collected from healthy adult animals ranging in ages from 7-23 years. To evaluate these methods for bias, we computed regressions between protein abundance and age for each protein with complete data. Then, missing values were introduced and imputed. Regression analysis was subsequently computed with the newly imputed results and the new regression coefficients and p-values were compared to truth. Unfortunately, MI methods often failed to converge as the number of proteins in the dataset approached 40, so for the comparison of MI and SI methods, either 10, 20, 30, or 40 proteins were randomly sampled from the complete data before introducing missing values and computing the imputation. Our results indicate that when randomly sampling 10 to 40 proteins and introducing missingness at rates between 10 to 40%, none of the imputation methods significantly biases the regression coefficients or p-values in either direction (Fig. 4). With a smaller number of proteins, however, there is suggestive evidence that LLS imputation may falsely inflate p-values (Fig. 4B). Conversely, MICE-Norm appears to be slightly conservative as the number of proteins grows (Fig. 4B). Of all the methods, Random Forest imputation appears just as unbiased, if not more unbiased, than any of the other imputation techniques – including the MI methods.

**Fig. 4.**
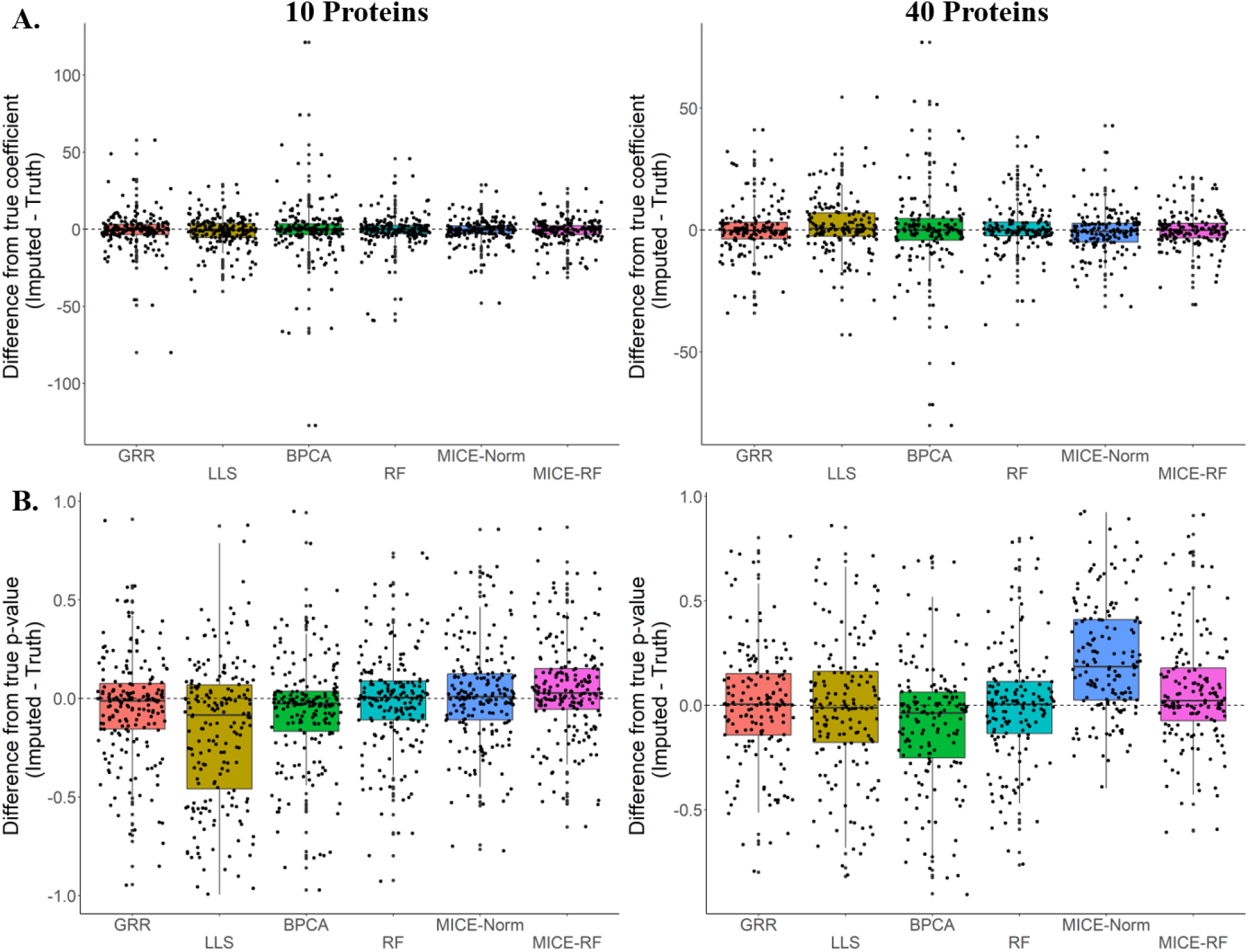
Differences between true regression estimates and imputed regression estimates (A.) as well as differences between association p-values (B.) for 6 imputation methods. Evaluations were done on randomly selected 10 and 40 proteins from complete data (296 proteins with no missing values). Global missingness of 10 to 40% was introduced for each individual protein list.

In order to more directly assess the influence of imputation on the inter-variable correlations, we computed Cronbach’s alpha (a measure of internal consistency) for a random subset of the complete data and compared it to the Cronbach’s alpha for the same subset of imputed data. Wherever the inter-variable correlation is decreased through imputation, Cronbach’s alpha will also decrease relative to the true data. Similarly, wherever the inter-variable correlation is increased through imputation, Cronbach’s alpha will increase. Any methods demonstrating consistent shifts in either direction away from the true Cronbach’s alpha may be biasing downstream results by altering the correlation structure between variables. This will be especially true for multivariate analyses. The suggestive patterns of bias seen among the differences in the regression coefficients and p-values were amplified in the Cronbach’s alpha analyses. LLS shows strong and significant bias in Cronbach’s alpha (Fig. 5). This bias appears to diminish as more proteins are added to the imputation. MICE-Norm shows suggestive evidence of a conservative bias for 20% global missingness and 30 randomly selected proteins. Random Forest demonstrates a slight, yet significant conservative bias for 20% global missingness and 30 randomly selected proteins.

**Fig. 5.**
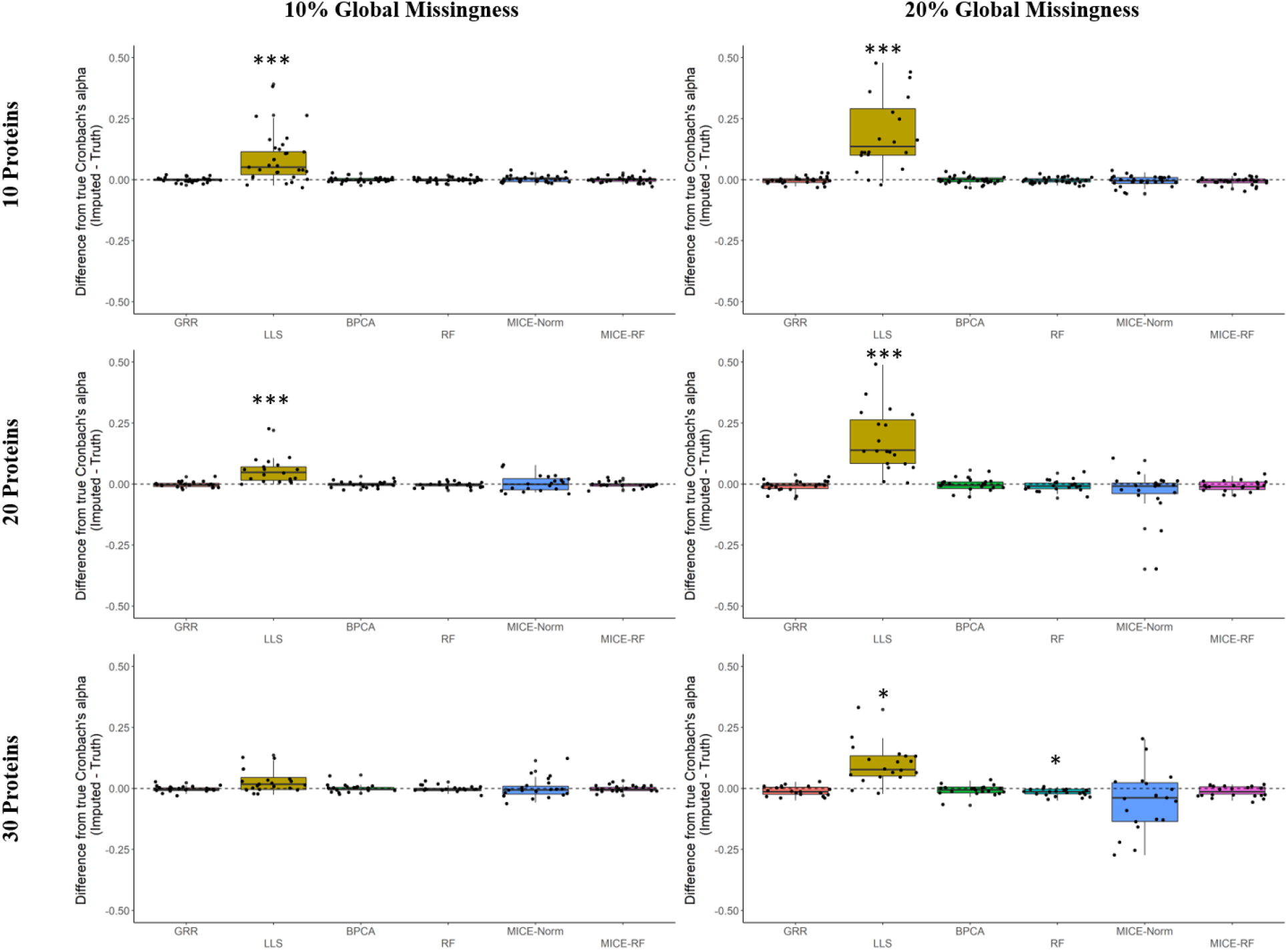
Differences between true Cronbach’s alpha and imputed Cronbach’s alpha for 6 imputation methods. (*** Bonferroni p < 0.001 * Bonferroni p< 0.05). Evaluations were done on randomly selected 10, 20 or 30 proteins (left axis) from complete data (296 proteins with no missing values). Global missingness (top axis) of 10 and 20% was introduced for each individual protein list.

It is known that MI reduces bias in statistical analysis by creating multiple versions of the imputed data where the imputed values in each of the imputed datasets are allowed to vary by a random component that reflects the level of confidence the prediction algorithm has in a particular imputed value [11-15, 26]. However, in the context of high-dimensional proteomics data, we demonstrate that MI may not be necessary (or beneficial) to reduce bias when imputing missing values with proteomics data for association analysis. Our results even suggest that MICE-Norm (the parametric MI technique) is overestimating the uncertainty in the imputed data and incorporating too much noise as the number of variables grows (Fig. 4 & 5). A direct example of what happens with the imputed values from MICE-Norm demonstrates how the variance quickly balloons when the number of variables grows large (Supp. Fig. 3). It is worth noting, however, that this problem is not evident in the non-parametric approach (Supp. Fig. 4). In addition to these results, we have found that MI often fails to converge with a large number of variables. Therefore, advanced variable selection would need to be computed in order for MI methods to work with high-dimensional proteomics data [9, 10]. MI also greatly complicates potential multi-omic studies – particular where users are interested in directly testing joint hypotheses across the multiple data types. Even though MI may ultimately protect from additional false positives, we believe that proceeding with SI is a very sensible approach.

Among the SI techniques: Random forest, GRR, and BPCA are all reasonable imputation approaches for the analysis of our data. They each show high amounts of accuracy, and little bias. Random Forest is a logical choice for imputation and has been recommended in the literature multiple times as a good imputation method for proteomics data and metabolomics data [7, 27-29]. In addition, Random Forest benefits from being a non-parametric approach, which is critical in data that is not always normally distributed. It is also less sensitive to outliers and noise [25]. The Random Forest imputation algorithm (missForest) that is implemented in this study first fills in missing values with the row medians, after marking each row that contains missing values as the prediction set and the complete data as the training set [25]. Then it begins an iterative process of looping through missing data points several times, using the newly updated data to inform the previously missing data point. It repeats this until either the algorithm converges or the number of user-defined iterations elapse [25]. Because the MI methods limited the number of proteins and the amount of missingness that could be used to previously evaluate the Random Forest algorithm, we wanted to complete a more in-depth evaluation of the Random Forest algorithm across higher percentages of missingness and with all proteins. To approximate the point of protein missingness at which the accuracy of the Random Forest algorithm begins to deteriorate, we used the protein full dataset (all 1622 proteins filtered down to 1252 proteins that occurred in at least 25% of samples) instead of just the complete proteins. Here, we introduced one missing value at a time where there was not already a missing value, imputed it along with all of the other real missing data points, and compared it to the real value. This was done in order to evaluate the accuracy of the Random Forest imputation algorithm while maintaining the exact data structure of the real data. All the way up to proteins with more than 75% missing data, Random Forest does not show a significant drop in accuracy (Fig. 6A). We hypothesize that this may also be a function of our data, where the relative amounts of protein across each of the samples remain quite stable as the protein mean imputation method also demonstrated such high accuracy with these data (Fig. 3). The Cronbach’s alpha across all proteins after imputation with the Random Forest algorithm across all proteins is significantly inflated for all amounts of overall missingness until the overall missingness reaches 70% (Fig. 6B). This means that users need to be aware that implementing this imputation with proteomics data is likely to falsely inflate correlations between variables. Despite this, our results show no significant increase or decrease in the estimates of the regression coefficients and association p-values (Fig. 6C and 6D). It is worth noting, however, that as the proportion of overall missingness increases, the regression coefficients of the imputed data start to deviate more and more from truth (Fig. 6C). For this reason, imputing proteins with the Random Forest method when there is greater than 30% protein missingness and datasets with more than 20% of overall missingness is not advised.

**Fig. 6.**
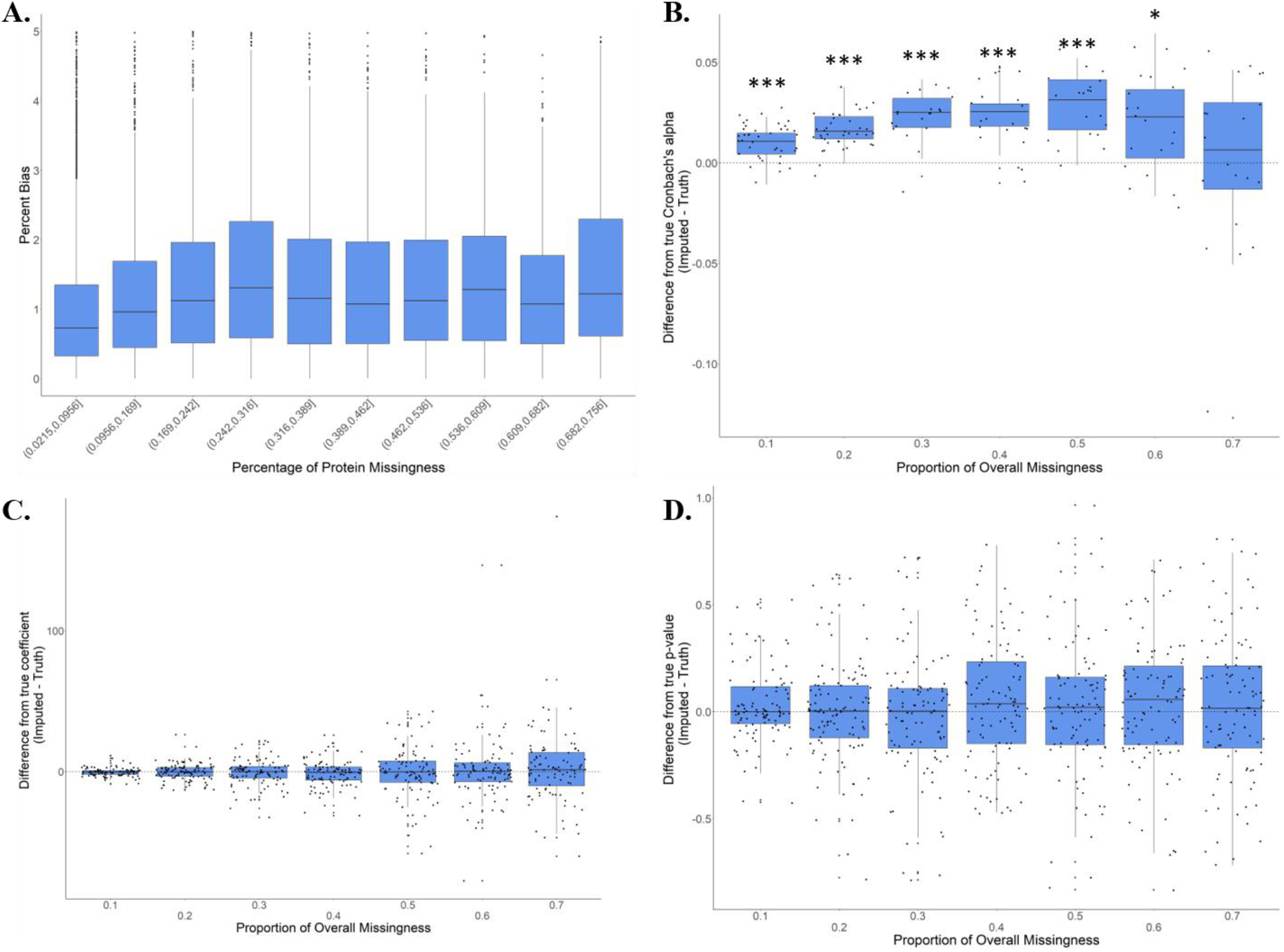
In depth Evaluation of Random forest imputation for the full protein dataset (proteins present in at-least 25% of samples, corresponding to 1252 proteins) (<u>*** Bonferroni p < 0.001 * Bonferroni p< 0.05</u>)

## Conclusions

Our current workflow for improving the efficiency of LFQ proteomics for NHPs by improved data search for enhanced PTM discovery and detailed evaluation of various methods for imputing the missing data can be very helpful for improving qualitative depth and quantitative accuracy of LFQ proteomic studies for NHPs. Our study incorporated the MetaMorpheus data search which improved the detection range of proteins including 58 different PTMs, and we report a number of novel PTMs. We also evaluated multiple imputation methods for improving the statistical analysis of proteomics data for NHPs, and we discussed the costs and benefits associated with MI in high-dimensional proteomics data. An implementation of the MetaMorpheus search workflow with an imputation of protein abundances for missing data with the Random Forest method is highly reliable when fewer than 30% of values for a particular protein (and less than 20% of datapoints overall) are missing. This standard LFQ workflow for NHP data also highlights the clinical translational opportunity in *H. sapiens* where discovery studies for control prefrontal cortex samples across a diverse age range is limited because of sample availability restrictions.

## Material and Methods

### Animal model

Our study included 45 *P. anubis* brain frontal cortex samples, consisting of 35 females in the age range of 7-23 years (21-69 human-equivalent years) and 10 males in the age range of 8-23 years (24-69 human equivalent years).

### Proteomics Sample Preparation

Samples were homogenized in Tris buffer. Tissue homogenate was subjected to an overnight Acetone precipitation at -20°C followed by centrifugation at 12,000g for 10 minutes. The protein pellet was dried and reconstituted in 100mM of Ammonium Bicarbonate. After protein quantification, 100ug of each sample was processed further. Protein disulfide bonds were reduced using dithiothriotol (DTT) for 1hour at 56°C followed by alkylation using iodoacetamide (IAA) for 30minutes in dark. The resulting protein mixture was subjected to an overnight trypsin digestion. Samples were desalted and cleaned up using Thermo Scientific Pierce C18 Tips. The final peptide mixture was dried and reconstituted in 0.1% formic acid.

### Data acquisition

1ug of each sample was loaded on a PepMap RSLC C18 easy-spray column (3um, 100A, 75um*15cm) using Easy-nLC 1200 coupled to an Orbitrap Lumos Tribrid Mass Spectrometer (Thermo Scientific). The peptides were separated using a 3-hour gradient of Mobile phase A (0.1% Formic acid in 95:5 Water: Acetonitrile) and Mobile Phase B (0.1% Formic acid in 80:20 Acetonitrile: Water). The following gradient program was used for peptide elution: 2 to 30% B in 140 minutes, 30% to 95% B in 30 minutes and 95-100%B in 10 minutes. The mass spectrometer data was acquired in MS1 scan mode (m/z=375-1800) at a resolution of 120,000 with Automatic Gain Control (AGC) of 4.0×10^5^ and Maximum injection time of 50 ms. MS/MS data acquisition was done using HCD mode at a resolution of 30,000 with an AGC target of 5.0×10^4^ and maximum injection time of 54 ms. All the data acquisition was done using Thermo Scientific Xcalibur software.

### Data Analysis

Raw data were analyzed using MetaMorpheus, using the *P. anubis* reference proteome database from Uniprot with 44,721 entries (UP000028761). The data files were calibrated using following settings: precursor mass tolerance of 15ppm, product mass tolerance of 25ppm with Carbamidomethyl as fixed modification and oxidation of methionine as variable modification. Trypsin was selected as protease with 2 maximum mixed cleavages and the calibrated data files were converted to mzML format. Post calibration data was searched using G-PTM-D task for incorporation of common biological, metal or artifact PTMs into the search database. Final search was done using the augmented search database with incorporated G-PTM-D based modifications at precursor and product mass tolerance of 5 and 20ppm respectively. Peptide or Protein quantification was done using the FlashLFQ approach and the Match between runs option was enabled. For both protein and peptide data files, only proteins or peptides identified in target database were retained. For PTM discovery, all peptide data were filtered with a q-value threshold of ≤0.01, precursor mass error of 5ppm and presence of at least 1 unambigous PSM (q<0.01). The protein data were normalized and log transformed using global intensity normalization. Only proteins present in at least 25% of the samples were retained for downstream imputation analysis.

### Introduction of Missing Values

For all of the initial imputation evaluations only the complete data (i.e., proteins with no missingness) were used. Missingness was introduced to the complete data at varying rates and with four different missing mechanisms – missing completely at random (MCAR), missing at random (MAR), missing not at random (MNAR), and mixture of all three (MCAR-MAR-MNAR), using approaches previously described [27]. Briefly, MCAR values were introduced by randomly selecting values to remove. MAR values were introduced by sorting the abundance values of two proteins using only one of the protein’s values. For the other protein not used to do the sorting, values were removed if the matched values (values shared across samples) in the protein used for sorting were above a cutpoint sampled from a chi-squared distribution. This is done to simulate a situation where high abundance in one protein leads to missingness in another protein. MNAR values were introduced by sampling a cut point from a chi-squared distribution and removing all values within the same protein below that cut point. For the mixture of all three missingness mechanisms, each missing type was sequentially added to the data at equal proportions until the desired rate of missingness was achieved. The mixture of all three missingness mechanisms (MCAR-MAR-MNAR) was considered most heavily out of all of the simulations because all three types of missingness are likely present in our data and influencing the imputation procedures.

### Evaluation of Imputation Methods

NAguideR was used beforehand to explore the various imputation methods that are commonly applied with proteomics data [6]. Thirteen SI methods (i.e., methods that predict the best value for each data point and treat it as truth in statistical analysis) were initially evaluated for accuracy. Percent bias (i.e., the percent difference between the imputed value and truth) was observed for each method and the percentage of data points with a percent bias <5% was also recorded and observed for each type of missingness and for varying degrees of protein missingness and global missingness. Four SI methods (GRR, LLS, BPCA, and Random Forest) were further evaluated for accuracy.

The most consistently accurate SI methods were also compared with MI methods (MICE-Norm and MICE-Random Forest). First, we compared the accuracy of regression coefficients from downstream association analysis between SI methods, truth, and the pooled results from MI methods. When the number of proteins in the data approached 40, imputation became very slow and computationally expensive. In addition, when the number of proteins in the data approached 50 and when there were high proportions of missingness, the MI methods frequently failed to converge and returned the following error: “system is computationally singular”. To circumvent this error, a random subset of either 10, 20, 30, or 40 proteins were sampled from the data. For the comparison of regression coefficients, we first introduced missing values into the complete data as described above and then sampled our random set of proteins before downstream analysis. These data were then merged with the available age information for each sample and, subsequently, the missing values were imputed. For the SI methods, a simple regression analysis was computed between age and protein abundance, treating the imputed values as truth. For the MI methods, a regression analysis was computed on each of the multiply imputed versions of the data and the results from each analysis were pooled according to Rubin’s rules to obtain a single estimate and p-value [14]. The differences between the regression coefficients and p-values from models using the complete data and regression coefficients from the models using the imputed data were recorded for each iteration of missingness that was introduced into the complete data. This was computed for global missingness rates of 10% and 20% as well as each of the numbers of randomly selected proteins, and each of the four missingness types. For each method, we computed a one sample t-test on the differences between the imputed and true regression coefficients and p-values to test for departure from zero.

We also compared the Cronbach’s alpha (a measure of internal consistency) between the imputed results and truth (i.e. the actual protein value that was removed from the original data). Once again, a random subset of either 10, 20, or 30 proteins were sampled from the complete data. The Cronbach’s alpha was computed with the complete data and then missing values were introduced and imputation completed. Cronbach’s alpha was again computed on the imputed results and compared with the true Cronbach’s alpha. For the MI methods, Cronbach’s alpha was computed on only 1 of the 5 multiply imputed datasets. The difference between the two measures was recorded and the process was repeated iteratively for global missingness rates of either 10% or 20% and each of the four missingness types. Higher numbers of proteins and higher rates of global missingness were not observed because the MI methods frequently failed to converge. A more in-depth evaluation of the Cronbach’s alpha was computed when MI methods were no longer being considered. For each method, we computed a one sample t-test on the differences between the imputed and true Cronbach’s alpha to test for departure from zero.

A more in-depth evaluation of accuracy and bias was carried out for the Random Forest method to determine the amount of missingness at which the method’s accuracy starts to deteriorate and where it begins to introduce biases. To maintain data structure as close to reality as possible, we used the full data after removing proteins with >75% missing data and randomly selected only a single non-missing data point at a time, set it to missing, and then imputed it. This was done iteratively for 10,000 of the possible 42,433 non-missing values present in the data. For each iteration, the true value was recorded along with the imputed value as well as the percentage of missing data in the protein being imputed.

In addition to these evaluations, we also computed further evaluations of the Cronbach’s alpha and regression coefficients for the Random Forest method. These were computed in an identical fashion to how they were computed for all methods, just without restrictions on the number of proteins used (which were in place to allow the MI methods to compute) and with a wider range of overall missingness rates (10% - 70%).

## Abbreviations

(MS): Mass Spectrometry
(LFQ): Label-free quantification
(PTMs): Posttranslational modifications
(RegImpute): Ridge Regression
(missForest): Random Forest
(LLS): local least squares
(BPCA): Bayesian Principal Component Analysis method
(CA3): Carbonic anhydrase 3
(MS/MS): Tandem Mass Spectrometry
(NHP): Non-human primates
(MI): Multiple Imputation
(SI): Single Imputation

## Declarations

### Ethics approval

All animal procedures were approved by the Texas Biomedical Research Institute (TBRI) Institutional Animal Care and Use Committee (IACUC) and conducted in Association for Assessment and Accreditation of Laboratory Animal Care (AAALAC) international-approved facilities.

### Availability of data and materials

The raw datasets used during the current study are available from the corresponding author on reasonable request.

### Competing interests

The authors declare that they have no competing interests

### Funding

This study was supported by NIH through grant U19 AG057758.

### Author’s contributions

ZH, KDZ and MO conceived the study. CL and PN overlooked animal experiments and provided all tissue samples. ZH and MO generated and analysed proteomics data. ZH, KDZ, HG and MO perfomed imputation analysis. ZH, KDZ and MO drafted the manuscript. HGA, LAC and PN provided significant editorial revisions.

## Acknowledgements

We want to acknowledge all the colleagues in our laboratory for useful suggestions and discussions. We are also thankful to the editor and reviewers for there careful reading and thoughtful comments.

## Supplementary Information

**Supp. File 1:** All protein and peptide list detected in our study (Protein intensity values are normalized to total ion intensity).

**Supp. File 2:** List of high confident PTMs detected in our study.

**Supp. File 3:** All peptides detected for reference protein carbonic anhydrase 3 (CA3). List of Modifications detected for peptide “DIKHDPSLQPWSVSYDGGSAK” from CA3.

**Supp. File 4:** Sequence coverage of reference protein carbonic anhydrase 3 (CA3) observed in our study and MS/MS spectral match of various PTMs identified for reference peptide from CA3.

**Supp. File 5:** List and brief description of all imputation methods used in this study.

## Supplementary figures

**Supp. Fig. 1:**
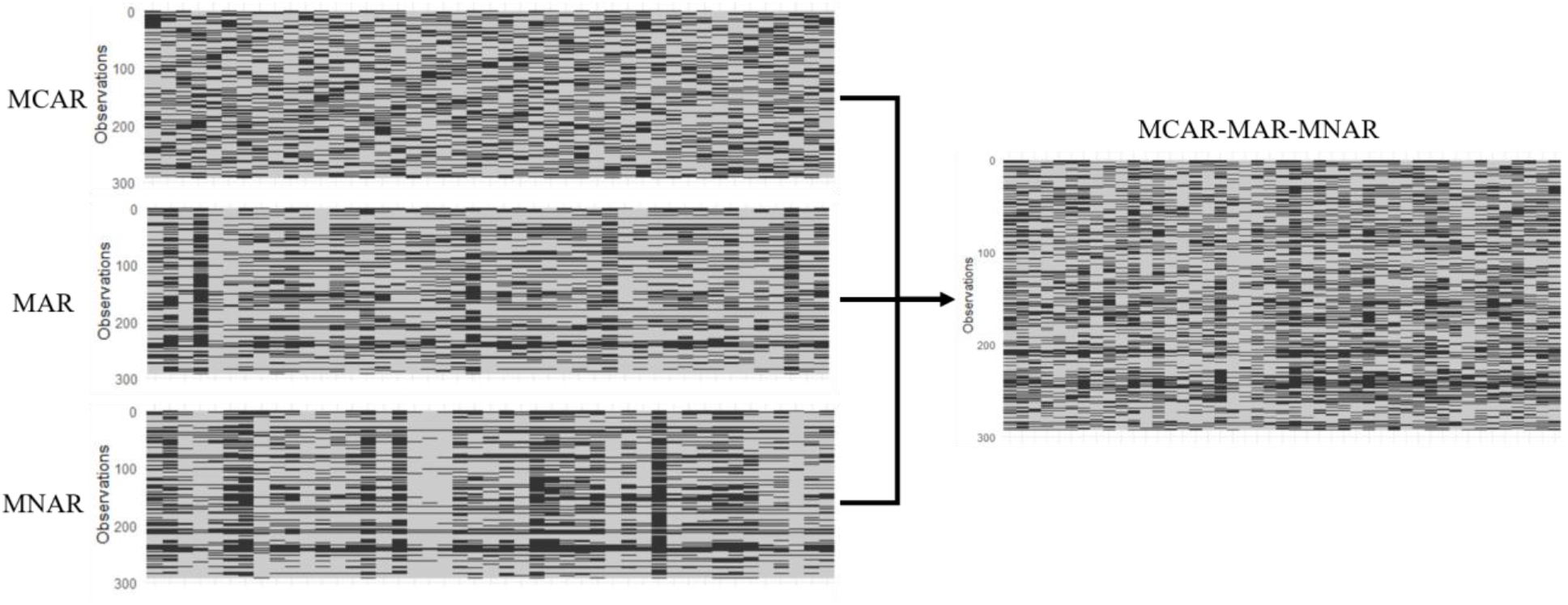
Introduction of different types of missingness in the protein list (The original list corresponds to 296 proteins with no missing values in the original dataset).

**Supp. Fig. 2:**
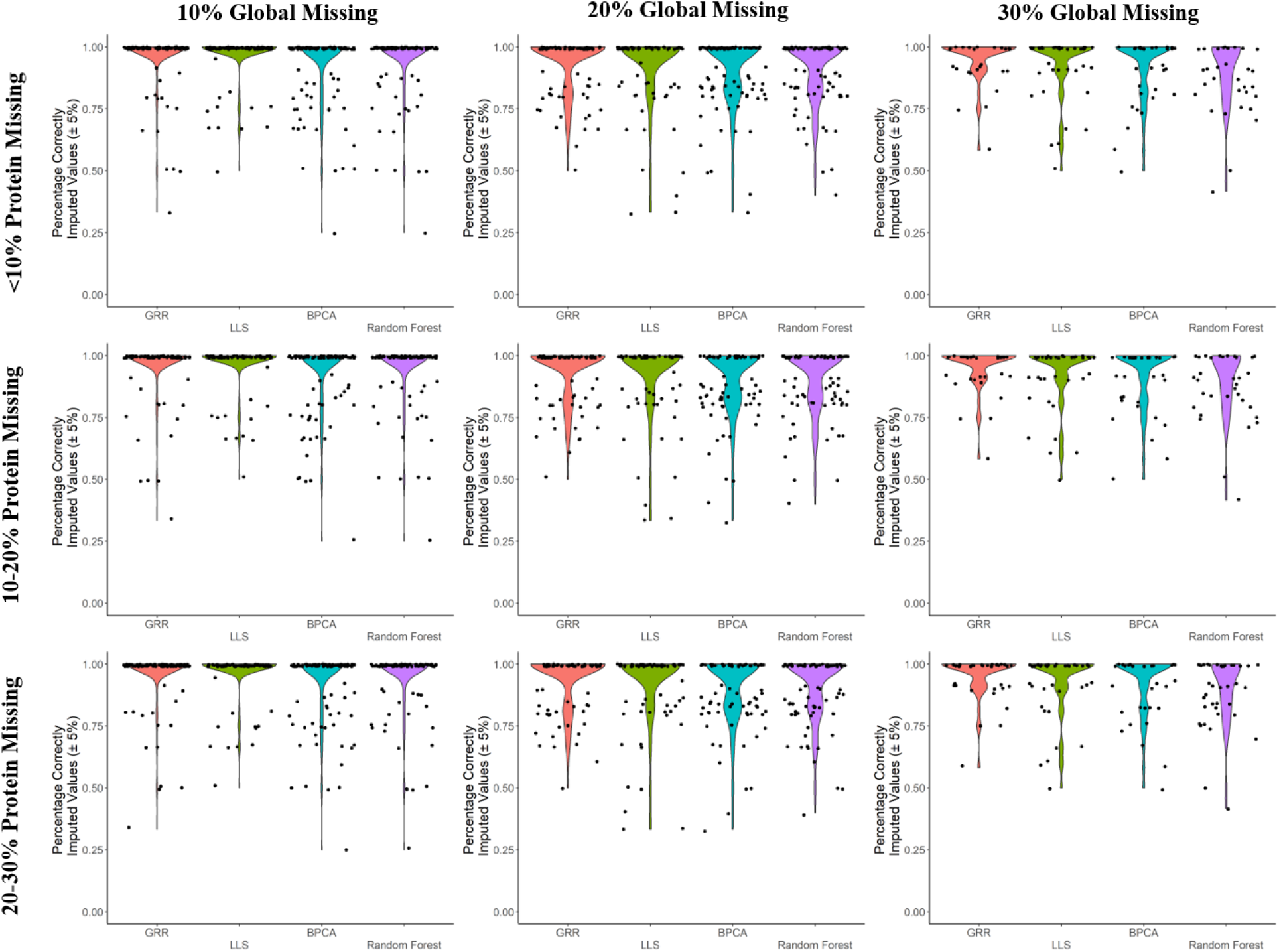
Evaluation of four single imputation methods (GRR, LLS, BPCA and Random Forest) at three different levels of missingness (<10%, 10-20% and 20-30%).

**Supp. Fig. 3:**
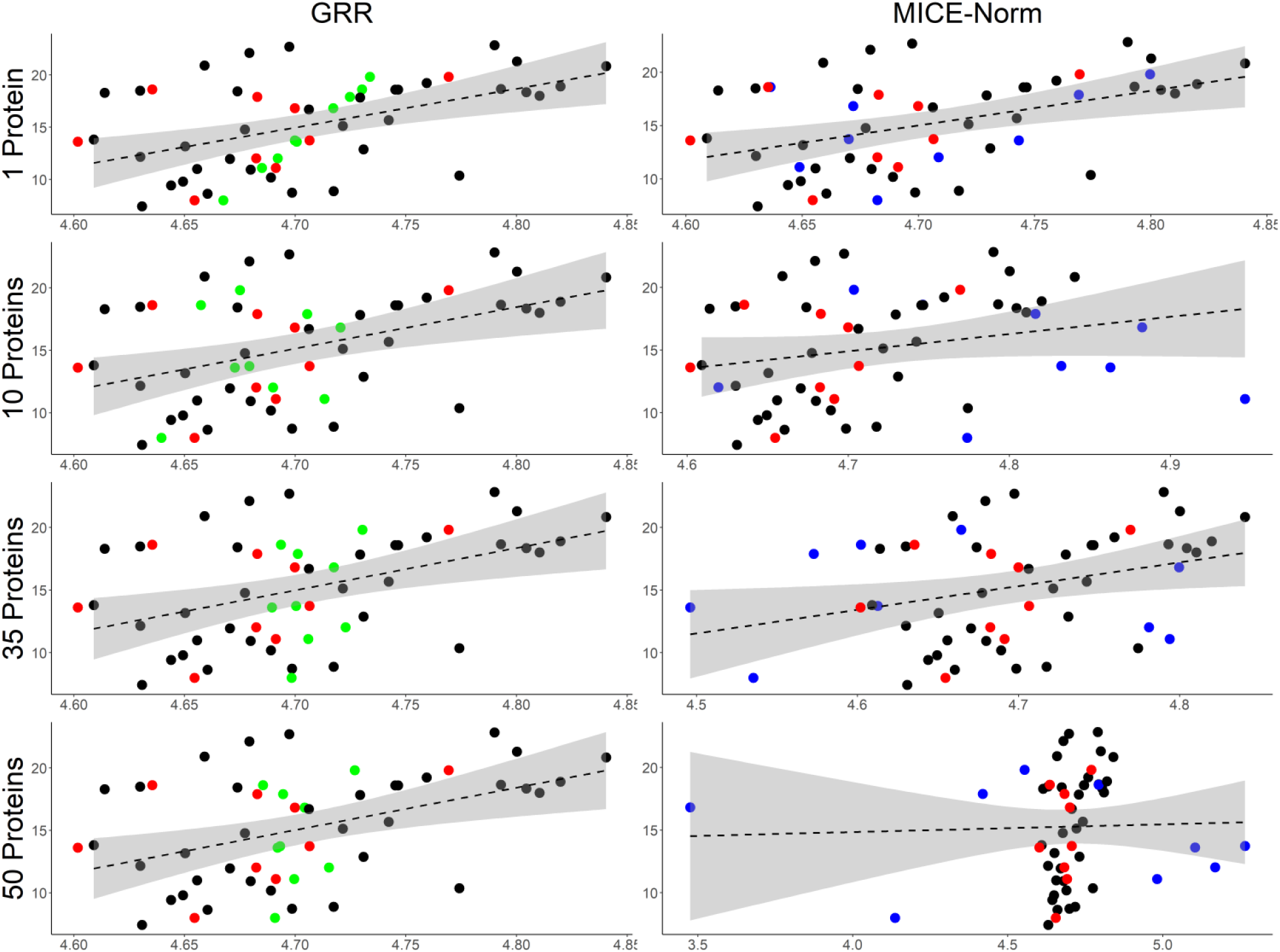
Example of the performance of imputation with parametric Single (GRR) and Multiple Imputation (MICE-Norm) methods. X-axis is protein intensity. Y-axis is “P. Anubis” age. Red is truth. Green are values imputed via GRR. Blue are values imputed via MICE-Norm.

**Supp. Fig. 4:**
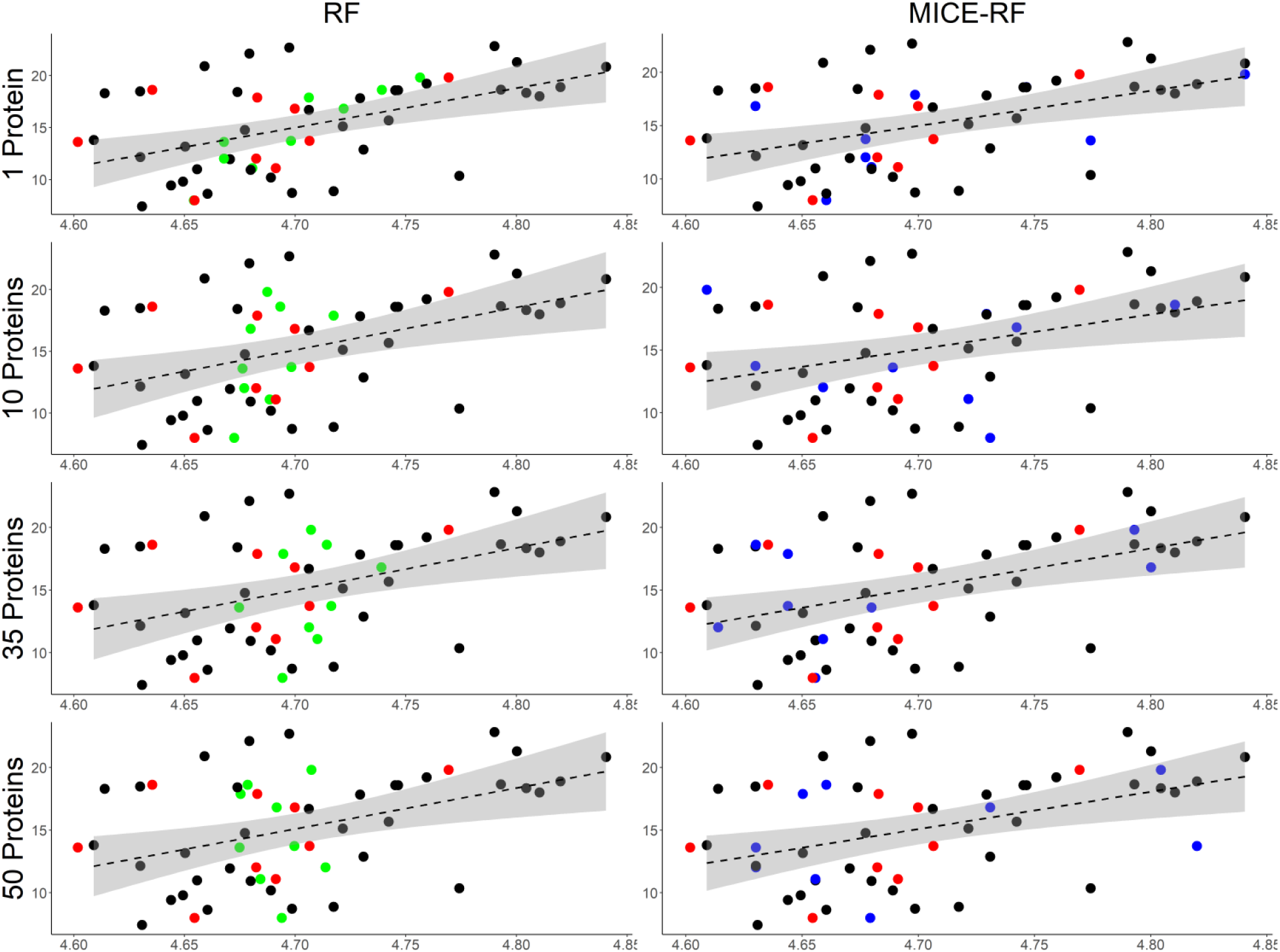
Example of the performance of imputation with non-parametric Single (RF) and Multiple Imputation (MICE-RF) methods. X-axis is protein intensity. Y-axis is “P. Anubis” age. Red is truth. Green are values imputed via RF. Blue are values imputed via MICE-RF.

## Notes

### Competing Interest Statement

The authors have declared no competing interest.

## References

1. Moulder R, Goo YA, Goodlett DR: Label-Free Quantitation for Clinical Proteomics. Methods Mol Biol 2016, 1410:65–76.

2. Filiou MD, Martins-de-Souza D, Guest PC, Bahn S, Turck CW: To label or not to label: applications of quantitative proteomics in neuroscience research. Proteomics 2012, 12(4-5):736–747.

3. Wang M, You J, Bemis KG, Tegeler TJ, Brown DP: Label-free mass spectrometry-based protein quantification technologies in proteomic analysis. Brief Funct Genomic Proteomic 2008, 7(5):329–339.

4. Proffitt JM, Glenn J, Cesnik AJ, Jadhav A, Shortreed MR, Smith LM, Kavanagh K, Cox LA, Olivier M: Proteomics in non-human primates: utilizing RNA-Seq data to improve protein identification by mass spectrometry in vervet monkeys. BMC Genomics 2017, 18(1):877.

5. Lazar C, Gatto L, Ferro M, Bruley C, Burger T: Accounting for the Multiple Natures of Missing Values in Label-Free Quantitative Proteomics Data Sets to Compare Imputation Strategies. J Proteome Res 2016, 15(4):1116–1125.

6. Wang S, Li W, Hu L, Cheng J, Yang H, Liu Y: NAguideR: performing and prioritizing missing value imputations for consistent bottom-up proteomic analyses. Nucleic Acids Res 2020, 48(14):e83.

7. Jin L, Bi Y, Hu C, Qu J, Shen S, Wang X, Tian Y: A comparative study of evaluating missing value imputation methods in label-free proteomics. Sci Rep 2021, 11(1):1760.

8. Liu M, Dongre A: Proper imputation of missing values in proteomics datasets for differential expression analysis. Brief Bioinform 2020.

9. Yin X, Levy D, Willinger C, Adourian A, Larson MG: Multiple imputation and analysis for high-dimensional incomplete proteomics data. Stat Med 2016, 35(8):1315–1326.

10. Gardner ML, Freitas MA: 2020.

13. Donders AR, van der Heijden GJ, Stijnen T, Moons KG: Review: a gentle introduction to imputation of missing values. J Clin Epidemiol 2006, 59(10):1087–1091.

14. van Ginkel JR, Linting M, Rippe RCA, van der Voort A: Rebutting Existing Misconceptions About Multiple Imputation as a Method for Handling Missing Data. J Pers Assess 2020, 102(3):297–308.

15. Rubin DB: Multiple Imputation After 18+ Years. Journal of the American Statistical Association 1996, 91(434):473–489.

16. Lodder P: To Impute or not Impute, That’s the Question. In: Advising on research methods: Selected topics 2013. Huizen: Johannes van Kessel Publishing; 2014.

17. Solntsev SK, Shortreed MR, Frey BL, Smith LM: Enhanced Global Post-translational Modification Discovery with MetaMorpheus. J Proteome Res 2018, 17(5):1844–1851.

18. Zhao Y, Long Q: Multiple imputation in the presence of high-dimensional data. Stat Methods Med Res 2016, 25(5):2021–2035.

19. Harju AK, Bootorabi F, Kuuslahti M, Supuran CT, Parkkila S: Carbonic anhydrase III: a neglected isozyme is stepping into the limelight. J Enzyme Inhib Med Chem 2013, 28(2):231–239.

20. Lundby A, Secher A, Lage K, Nordsborg NB, Dmytriyev A, Lundby C, Olsen JV: Quantitative maps of protein phosphorylation sites across 14 different rat organs and tissues. Nat Commun 2012, 3:876.

21. Bisen S, Kakhniashvili D, Johnson DL, Bukiya AN: Proteomic Analysis of Baboon Cerebral Artery Reveals Potential Pathways of Damage by Prenatal Alcohol Exposure. Mol Cell Proteomics 2019, 18(2):294–307.

22. Oba S, Sato MA, Takemasa I, Monden M, Matsubara K, Ishii S: A Bayesian missing value estimation method for gene expression profile data. Bioinformatics 2003, 19(16):2088–2096.

23. Troyanskaya O, Cantor M, Sherlock G, Brown P, Hastie T, Tibshirani R, Botstein D, Altman RB: Missing value estimation methods for DNA microarrays. Bioinformatics 2001, 17(6):520–525.

24. Ma W, Kim S, Chowdhury S, Li Z, Yang M, Yoo S, Petralia F, Jacobsen J, Li JJ, Ge X et al: 2021.

25. Stekhoven DJ, Buhlmann P: MissForest--non-parametric missing value imputation for mixed-type data. Bioinformatics 2012, 28(1):112–118.

26. Doove LL, Van Buuren S, Dusseldorp E: Recursive partitioning for missing data imputation in the presence of interaction effects. Computational Statistics & Data Analysis 2014, 72:92–104.

27. Kokla M, Virtanen J, Kolehmainen M, Paananen J, Hanhineva K: Random forest-based imputation outperforms other methods for imputing LC-MS metabolomics data: a comparative study. BMC Bioinformatics 2019, 20(1):492.

28. Wei R, Wang J, Su M, Jia E, Chen S, Chen T, Ni Y: Missing Value Imputation Approach for Mass Spectrometry-based Metabolomics Data. Sci Rep 2018, 8(1):663.

