## Supplementary material for "Assessment of label-free quantification and missing value imputation for proteomics in non-human primates": Supp. File 4

**Panel A**: Sequence coverage of CA3 detected in our study is marked with bold alphabets in purple color corresponding to >90% of coverage. PTMs of underlined peptide "DIKHDPSLQPWSVSYDGGSAK” are described in more detail in panel B.

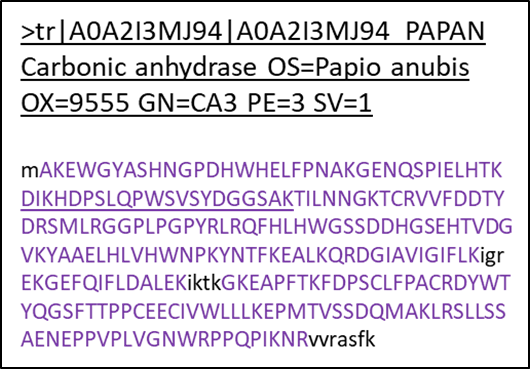

**Panel B**: Various modifications of underlined peptide "DIKHDPSLQPWSVSYDGGSAK” and corresponding MS/MS spectra match with “b” and “y” ions are shown below.

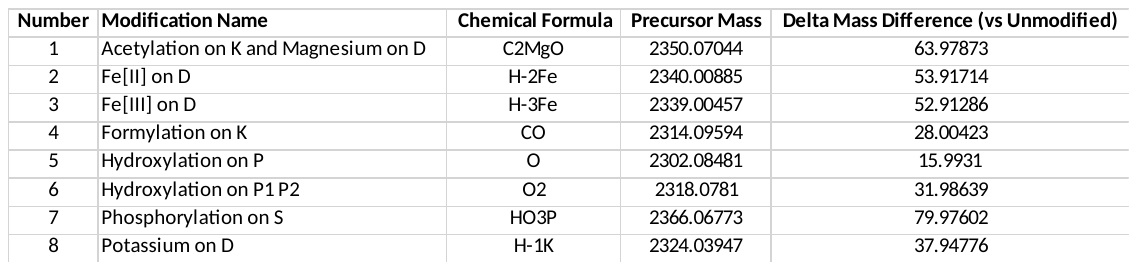

1) Acetylation on K and Magnesium on D

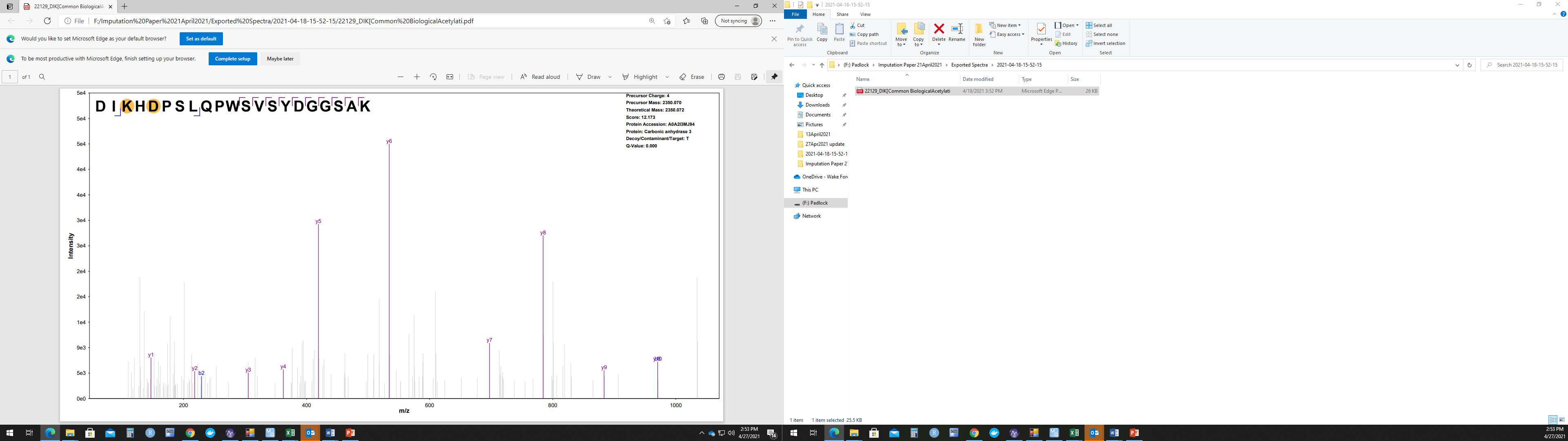

2) **Fe[II] on D**

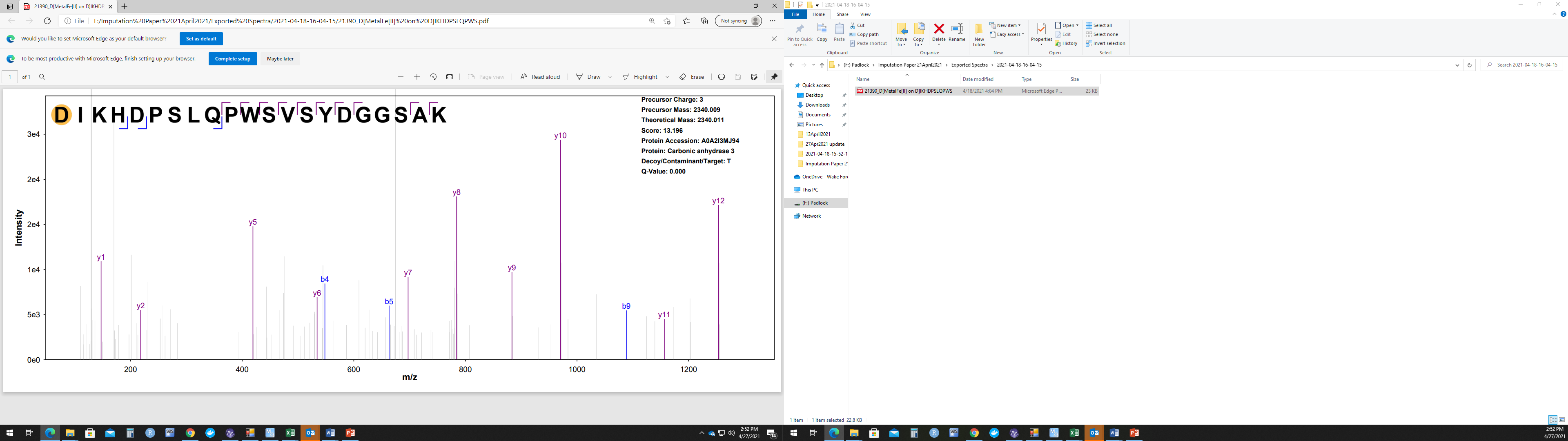

3) **Fe[III] on D**

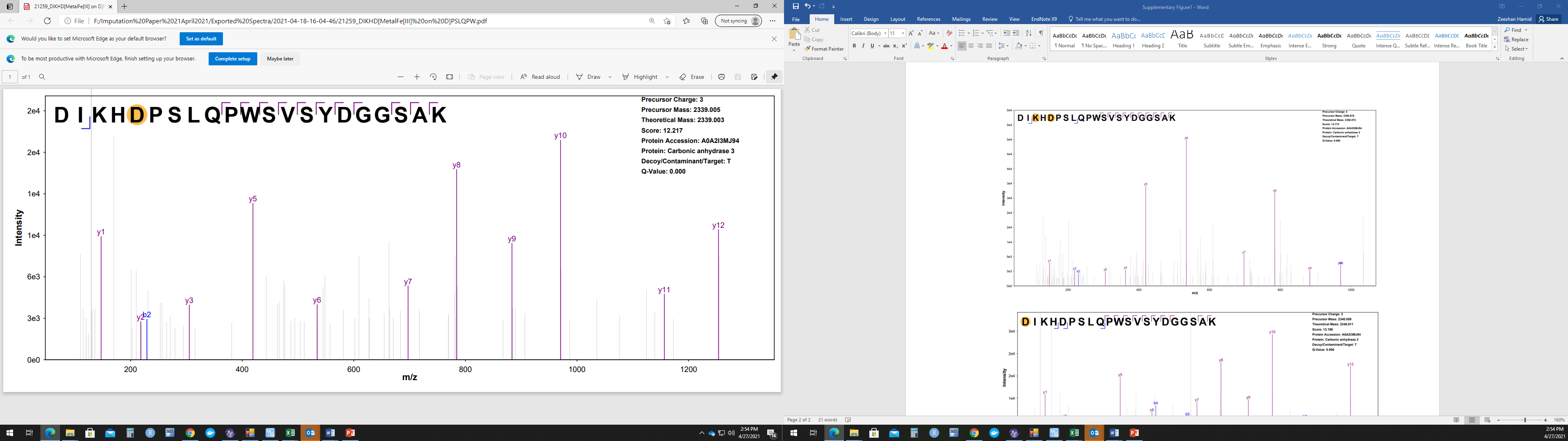

4) **Formylation on K**

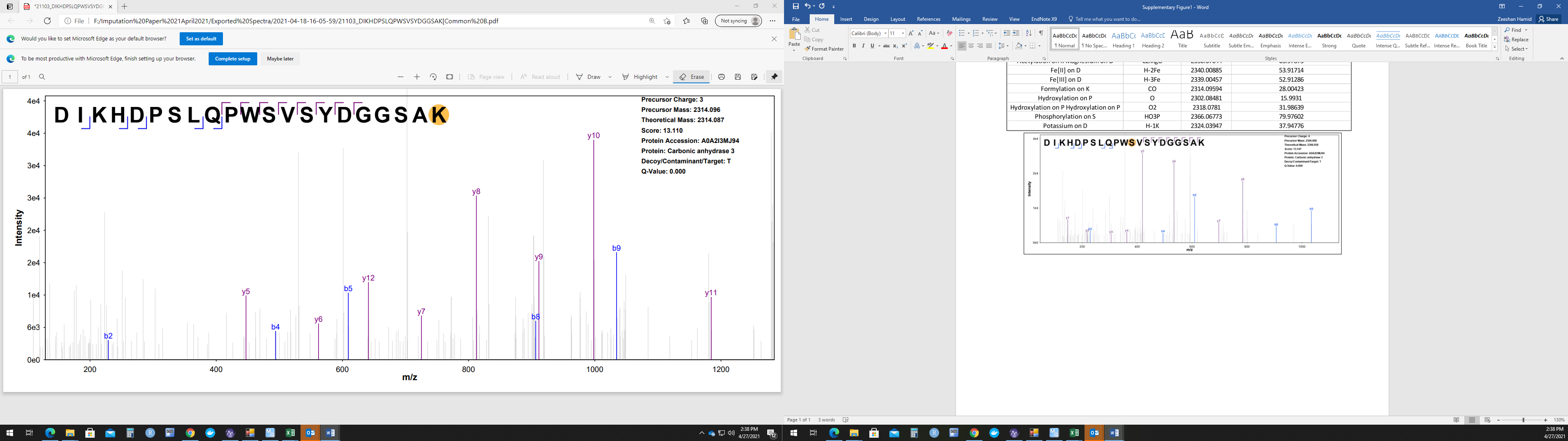

5) **Hydroxylation on P**

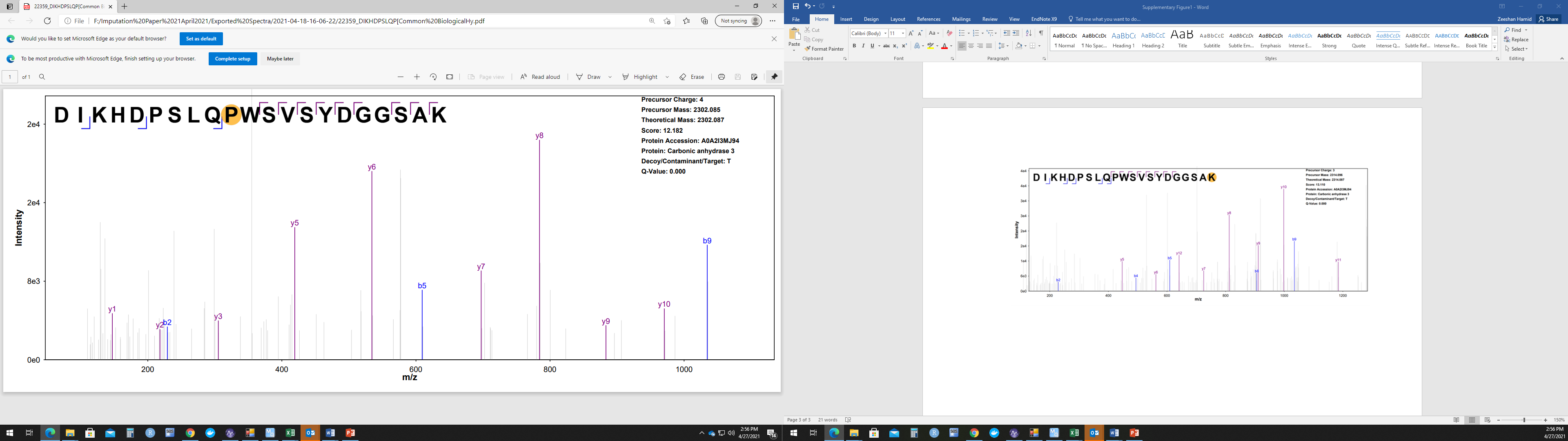

6) **Hydroxylation on P1 P2**

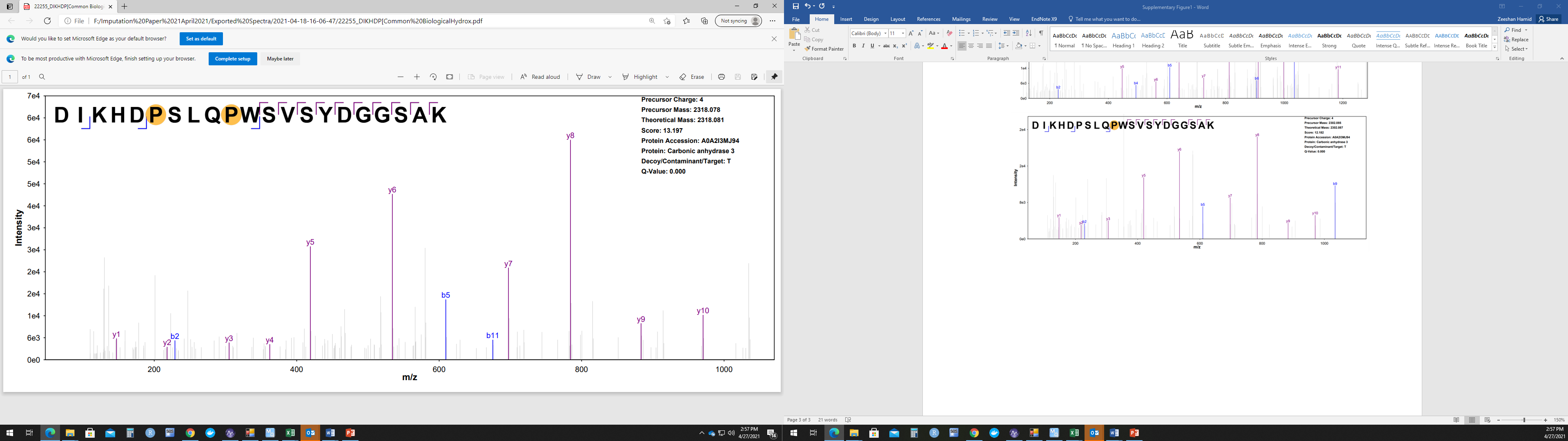

7) **Phosphorylation on S**

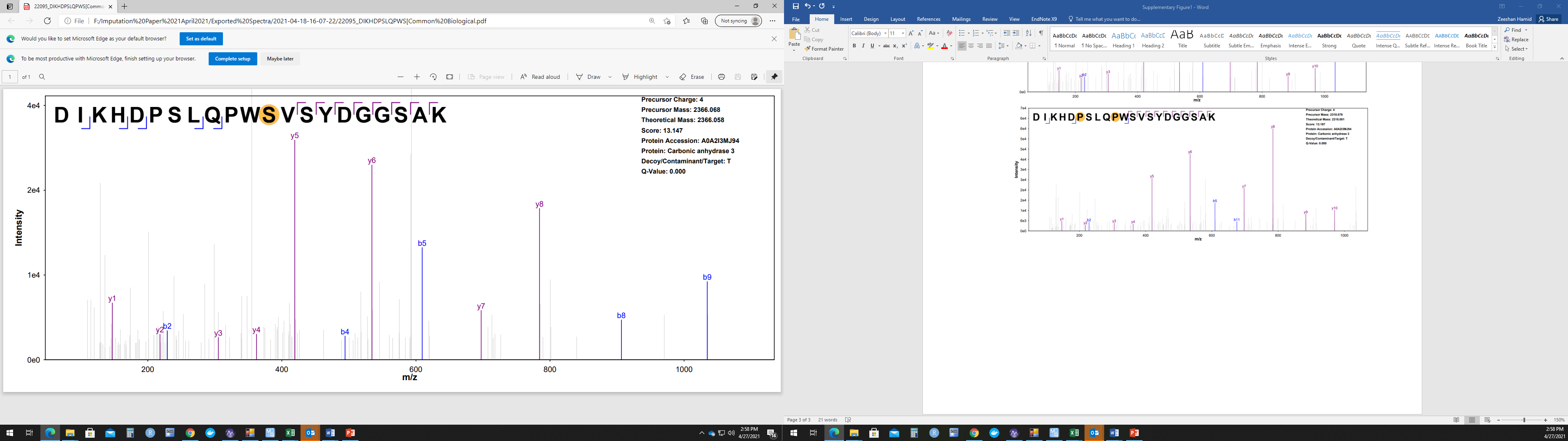

8) **Potassium on D**

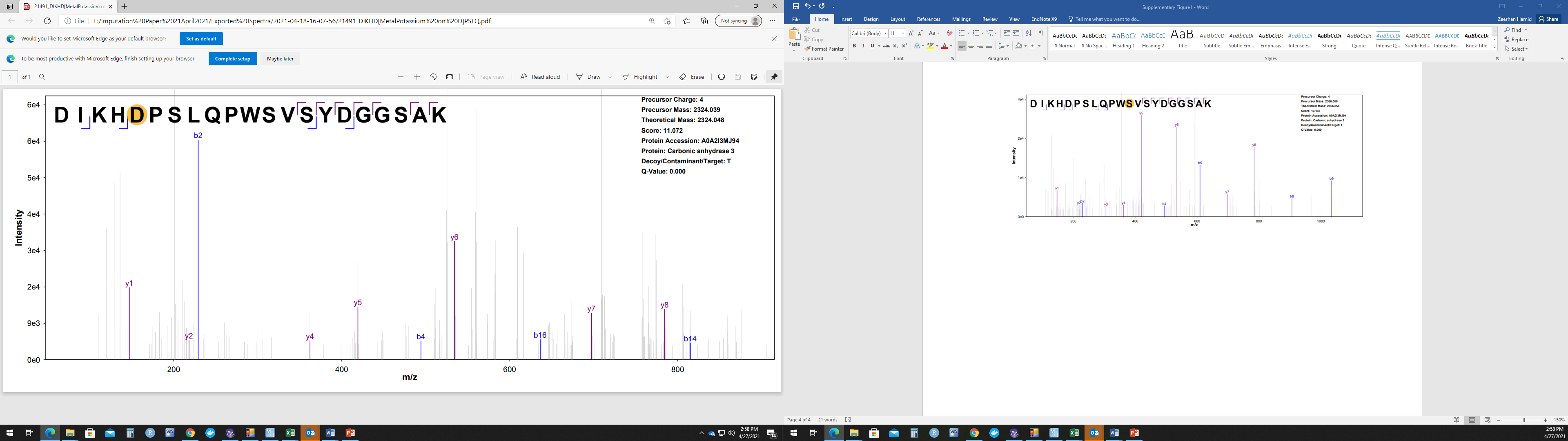
