## Supplementary material for "Assessment of label-free quantification and missing value imputation for proteomics in non-human primates": Supp. File 5

| **Imputation Method** | **Description** |
| --- | --- |
| **Single Imputation Methods** | |
| Glmnet Ridge regression (GRR) | This method is a regression based imputation method that uses an iterative procedure that refits multivariate prediction models that leverage the imputed values from the most recent iteration. This iterative procedure helps improve the prediction accuracy, and usually converges after 10 iterations |
| Local least squares(LLS) | This method was initially introduced for missing value imputation of microarray expression data. The way it is applied here, a target protein is represented as linear combination of any number of k similar proteins. Protein similarity is computed with Euclidean distance or Pearson correlation coefficients. The k selected variables are then are then used to predict the missing value of a target protein. (Kim, Golub et al. 2005) |
| Bayesian Principle Component Analysis (BPCA) | This method uses a Bayesian estimation to fit a probabilistic principal component analysis and missing value imputation using iterative expectation maximization algorithm. (Oba, Sato et al. 2003) |
| Random Forest(RF) | This imputation procedure implements a prediction model that is primarily based on non-missing values. In subsequent stepsit marks each row containing missing values as a part of the prediction set and each row with complete data as the training set. The program initially fills in missing values with row medians, but then iteratively fills in missing values and imputes to help improve prediction accuracy. (Stekhoven and Buhlmann 2012) |
| Mean | This method simply fills in missing values with the mean of the set of non-missing values. |
| KNN (k-nearest neighbor) | This method selects proteins with expression profiles similar to the target protein based on a Euclidean metric and imputes missing values by averaging the non-missing values of its nearest neighbors.(Troyanskaya, Cantor et al. 2001) |
| Sequential K-Nearest Neighbor (SeqKNN) | Modified version of the KNN method above in which missing values are imputed sequentially based on the proteins with least missing rate using a weighted mean of k nearest neighbors. (Kim, Kim et al. 2004) |
| ImpSeq | This method imputes missing multivariate data using a sequential algorithm. Each missing value is estimated by finding the value that minimizes the determinant of the covariance in data matrix. This value is then added to the complete data matrix and the algorithm continues to the next missing value. (Verboven, Branden et al. 2007) |
| SVD (Singular value decomposition) | This method employs singular value decomposition to obtain similar protein patterns that can be linearly combined to approximate the expression of all proteins in the data. Imputation using this method will initialize all missing elements with zero, followed by their estimation as a linear combination of the k most significant eigen-variables iteratively until reaches certain convergence threshold. (Troyanskaya, Cantor et al. 2001) |
| QR (Quantile regression) | This method uses random draws from the truncated distribution to perform imputation of left censored missing data. Parameters are estimated using quantile regression.(Wang, Li et al. 2020) |
| Maximum  likelihood  estimation (MLE) | This method used EMemploys maximum likelihood estimation with the EM algorithm to predict missing values from the global covariance structure among all samples (Wang, Li et al. 2020) |
| Zero | All missing values are replaced with zero |
| **Multiple Imputation Methods** (Yin, Levy et al. 2016, Wang, Li et al. 2020) |  |
| Mice cart | Multivariate Imputation by Chained Equations: classification and regression tress. This method generates multiple imputations for incomplete multivariate data by Bayesian linear regression |
| Mice norm | Multivariate Imputation by Chained Equations: Bayesian linear regression. This method generates multiple imputations for incomplete multivariate data by the random forest imputation algorithm described above. |

**References:**

Kim, K. Y., B. J. Kim and G. S. Yi (2004). "Reuse of imputed data in microarray analysis increases imputation efficiency." BMC Bioinformatics **5**: 160.

Oba, S., M. A. Sato, I. Takemasa, M. Monden, K. Matsubara and S. Ishii (2003). "A Bayesian missing value estimation method for gene expression profile data." Bioinformatics **19**(16): 2088-2096.

Stekhoven, D. J. and P. Buhlmann (2012). "MissForest--non-parametric missing value imputation for mixed-type data." Bioinformatics **28**(1): 112-118.

Troyanskaya, O., M. Cantor, G. Sherlock, P. Brown, T. Hastie, R. Tibshirani, D. Botstein and R. B. Altman (2001). "Missing value estimation methods for DNA microarrays." Bioinformatics **17**(6): 520-525.

Verboven, S., K. V. Branden and P. Goos (2007). "Sequential imputation for missing values." Comput Biol Chem **31**(5-6): 320-327.

Wang, S., W. Li, L. Hu, J. Cheng, H. Yang and Y. Liu (2020). "NAguideR: performing and prioritizing missing value imputations for consistent bottom-up proteomic analyses." Nucleic Acids Res **48**(14): e83.

Yin, X., D. Levy, C. Willinger, A. Adourian and M. G. Larson (2016). "Multiple imputation and analysis for high-dimensional incomplete proteomics data." Stat Med **35**(8): 1315-1326.
